# The cytoprotective role of antioxidants in mammalian cells under rapidly varying temperature, pressure and UV conditions during stratospheric balloon campaign

**DOI:** 10.1101/526376

**Authors:** Dawid Przystupski, Agata Górska, Paulina Rozborska, Weronika Bartosik, Olga Michel, Joanna Rossowska, Anna Szewczyk, Małgorzata Drąg-Zalesińska, Paulina Kasperkiewicz, Jędrzej Górski, Julita Kulbacka

## Abstract

Currently ongoing age of the dynamic development of the space industry brings the mankind closer to the routine manned space flights and space tourism. That progress leads to a demand for intensive astrobiological research aimed at improving strategies of the pharmacological protection of the human cells against extreme conditions. Although routine research in space remain out of our reach, it is worth noticing that unique severe environment of the Earth’s stratosphere have been found to mimic subcosmic conditions, giving rise to the opportunity for use of stratospheric surface as a research model for the astrobiological studies. Our study included launching balloon into the stratosphere containing the human normal and cancer cells treated with various compounds to examine whether these medicines are capable to protect the cells against the stress caused by rapidly varying temperature, pressure and radiation, especially UV. Due to oxidative stress caused by irradiation and temperature shock, we used natural compounds which display antioxidant properties, namely catechin isolated from green tea, honokiol derived from magnolia, curcumin from turmeric and cinnamon extract. “After-flight” laboratory tests displayed the most active antioxidants as potential agents which can minimize harmful impact of extreme conditions to the human cells.

## 2. Introduction

The world’s first hydrogen balloon was created over 200 years ago in Paris, which started the era of scientific ballooning(1). Dynamic development of this technique has enabled the advanced research in atmospheric science, aerobiology, and meteorology (2), remaining promising tool in astrobiology and space biology. In stratosphere the temperature generally drops down −40°C, atmospheric pressure is at 1 kPa, the relative humidity of air is lower than 1%, solar UV irradiance is about 100 W/m^2^ and cosmic radiation is at the level of 0.1 mGy/d ((3–5)). Stratospheric flights could potentially provide us with a valuable information about the stress response in living organisms after exposure to different severe environmental factors rapidly varying at the same time, which is hard to be mimicked in the laboratory. Moreover, we can examine if some medicines are able to support the viability of living organisms and cells in such an extreme environment. The exposure of varied biological samples to the stratospheric conditions opens the possibility to observe many changes in the cells’ functioning e.g. decreased viability, dysfunction of cellular organelles and their localization, cell cycle arrest, changes of gene expression and DNA damage (6,7). Additionally, irradiation in the stratosphere affects the cells either by direct or indirect pathways (water radiolysis, therefore exacerbating the oxidative stress (8)), causing e.g. mitochondrial function disturbance (9), DNA damage, proteins and lipids peroxidation correlated with disruption of the cell membrane (10), that altogether may lead to the cell death (8,10) (Fig 1). Furthermore, stratospheric flights provide unique cyclic changes of linked environmental factors, including radiation, overload, pressure, temperature, wind and vibrations, which are impossible to be simulated altogether in laboratory. The wide use of balloons provides numerous advantages including lower overall project costs, recoverable and massive payloads (up to 3600 kg) and more rapid and flexible flight(3).

**Fig 1.**
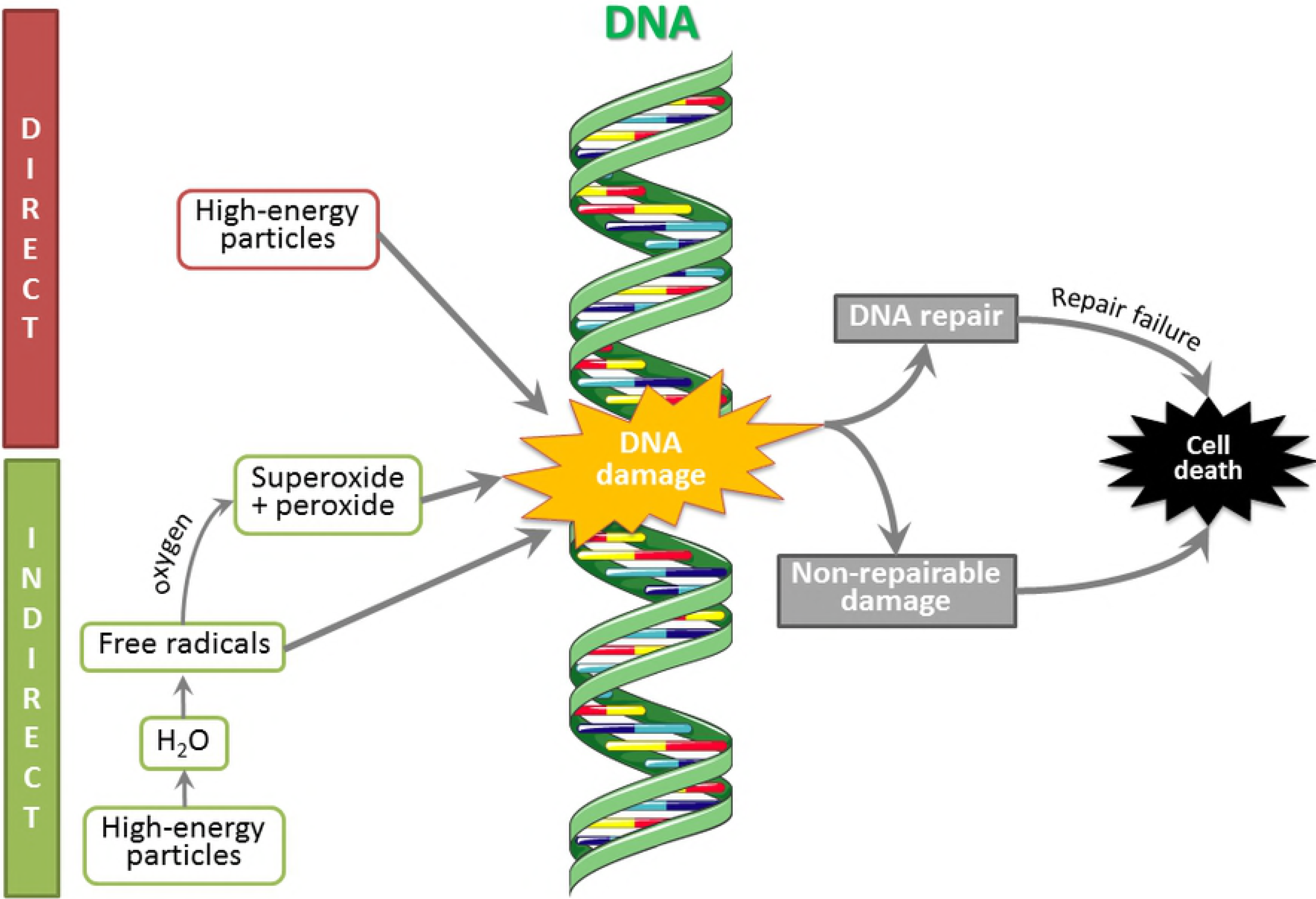
Effect of the radiation on the DNA. High-energy particles could affect DNA damage either by direct pathway or indirect pathway. Free radicals and by-products of the process of water radiolysis contribute to an increase of the oxidative stress in the cell affecting the DNA. Survival of the cell depends on the extent of the DNA damage (8–10). This figure was prepared using Servier Medical Art, available from www.servier.com/Powerpoint-image-bank.

It is known that natural substances derived from plants demonstrated positive impact on living cells including their protective role against diverse factors generating oxidative stress. Green tea polyphenols, such as epigallocatechin gallate, epicatechin gallate, epigallocatechin, epicatechin and catechin (11), are one of the strongest antioxidants and the long-term consumption of green tea extracts increases the activity of oxidative stress enzymes (12) (e.g. superoxide dismutase), inhibits lipo- and cyclooxygenase, xanthine oxidase, activator protein 1 and NF-κB transcription factors activity (13) and protects cells against radiation(14). Previous meta-analyses suggest a protective role of green tea against numerous cancer types (15–20). Similar properties are displayed by honokiol. This magnolia-derived compound inhibits UVB-induced immunosuppression and induces apoptosis in malignant cells (21). Chilampalli et al. (2010) showed that pretreatment with honokiol is efficient in preventing skin carcinogenesis induced by UV (22). Furthermore, honokiol inhibits photocarcinogenesis by targeting UVB-induced inflammatory mediators and cell cycle regulator (23,24). Turmeric compounds are another free radicals’ scavengers. A yellow spice curcumin displays anti-inflammatory (25), antibacterial (26), antiviral (27), anti-cancer (28), proapoptotic (29), neuroprotective (30,31), hepatoprotective (32) activity, promotes autophagy (33), affects the cytochrome P-450 enzyme system (34) and phase II enzymes (35). Curcumin is able to deactivate different forms of free radicals, such as reactive oxygen and nitrogen species (36), modulate the activity of GSH, catalase, and SOD enzymes which reduce oxidative stress (37,38). It can inhibit ROS-generating enzymes such as lipoxygenase/cyclooxygenase and xanthine hydrogenase/oxidase (39–41). Additionally, curcumin shows photosensitizing properties - it can absorb radiation of the appropriate wavelength damaging the tissue by free radicals and facilitating cell death (42). Different forms of free radicals can be scavenged by turmeric compounds. Similar properties are displayed by organic compounds found in cinnamon extract. Beside the antioxidant activity, cinnamon presents radioprotective potential. Azab et al. (2011) have revealed that extract of cinnamon improves the disturbance of the antioxidant system. Furthermore, this substance triggers protective action against protein and lipid oxidation via alteration of membrane structure (43). Thus, the main aim of this study was to evaluate the effect of stratospheric environment on two living cell lines: human ovarian cancer cells (SKOV-3) and Chinese hamster ovary cells (CHO-K1) in the presence of various natural compounds and investigate the protective role of these drugs. Additionally, the effect of radiation on the examined cells was observed. First, the adherent cells were seeded at equal density (500 000 cells/well) on 6-well plates. After 24-hour incubation with various antioxidant the cells were detached, suspended in freezing medium Bambanker™ (Nippon Genetics, Cat. no. BB01) (1,5 × 10^6^ cells/300 μL) and placed in microtubes 30 minutes before the balloon flight. Then the samples were transported on ice to the starting point and placed in a radiation transmitting gondola, located on the upper side of environmental measurement unit with accelerometer and temperature, pressure and UV sensors. One half of the samples was covered with aluminum foil to protect the cells against irradiation - mostly UV (the samples described in the study as “protected against radiation”), another half was sent into the stratosphere without the protective layer (described as “not protected against radiation”). As a result, we were able to evaluate the effect of radiation on examined cells in the presence of various antioxidants. As a control we used the appropriate number of cells incubated at 37°C in a humidified incubator with 5% CO2. Directly after landing, the biological samples were transported on ice to the specialized laboratory, where after-light tests were performed. The cells were seeded on 96-well plates (10 000 cells/well) and incubated in an appropriate drug solution for 24 hours to perform membrane permeabilization assay and intracellular reactive oxygen species generation assay, and for 24, 48 and 72 hours to evaluate mitochondrial activity in MTT assay. Modified versions of these assays (adding reagents into the cells’ suspension, appropriate incubation, centrifugation and seeding on 96-well plates) were used to analyze the suspended (not adherent to multiwell plates) cells directly after the balloon landing (0 h). Additionally, the cells were plated on 6-well plates and incubated in the appropriate drug solution for 24 hours to perform cell cycle assay, and for 7 days to carry out clonogenic assay. To analyze the expression of manganese-dependent superoxide dismutase (SOD2) the cells were seeded on 10-well diagnostic microscopic slides and fixed after 24 hours. Immunocytochemical staining was carried out in the two following days. Furthermore, neutral comet assay was performed to analyze DNA damages associated with the balloon flight. The scheme of the completed experiment is provided below (Fig 2).

**Fig 2.**
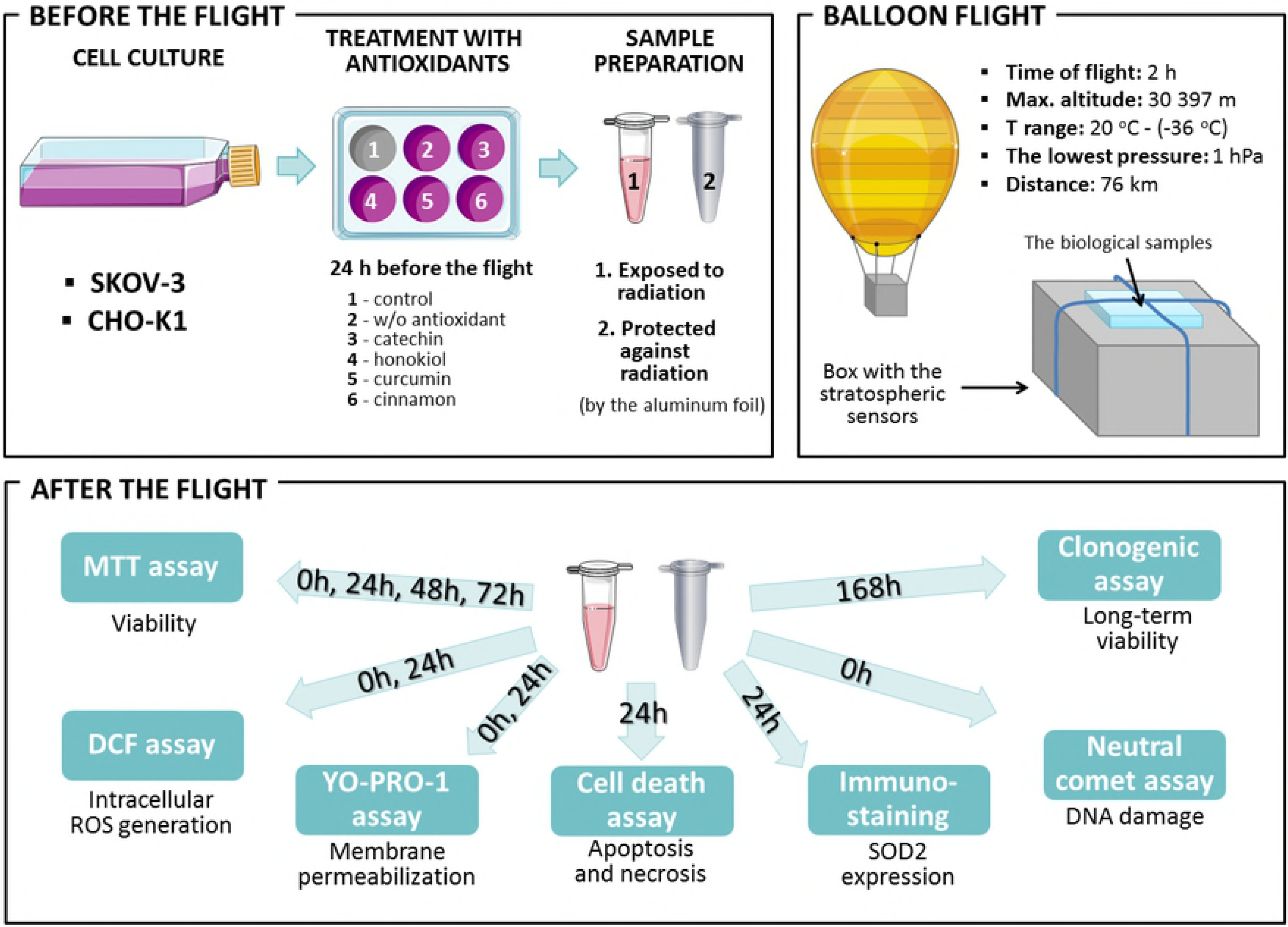
The schematic representation of the procedure of balloon flight and the preparation of biological samples. This figure was prepared using Servier Medical Art, available from www.servier.com/Powerpoint-image-bank.

## 3. Material and Methods

### 3.1. Cell culture

Human ovarian cancer cells (SKOV-3) and Chinese hamster ovary cells (CHO-K1) were obtained from the American Type Culture Collection (ATCC, London, UK). Cells were cultured as a monolayer in Dulbecco’s modified Eagle medium (DMEM, Sigma-Aldrich, USA; for SKOV-3 cells) and Ham’s F-10 Nutrient Mix (F-10, Sigma-Aldrich, USA; for CHO-K1 cells) containing 2 mM L-glutamine, 10% fetal bovine serum (FBS, Sigma-Aldrich) and 20 units penicillin and 20 µg streptomycin/mL (Sigma-Aldrich) at 37°C in a humidified incubator with 5% CO2. For the experiments, the cells were washed with Dulbecco’s Phosphate Buffered Saline (PBS, Bioshop, UK) and detached from the flask’s surface using 0.25% trypsin with 0.02% EDTA (Sigma-Aldrich, Poland).

### 3.2. Chemical substances

For the experiments, we used four compounds with antioxidative potential: (±)-catechin hydrate (Sigma, Cat. no. C1788), curcumin (Sigma-Aldrich, Cat. no. C1386), honokiol (Sigma-Aldrich, Cat. no. H4914) and cinnamon oil (Avicenna-Oil, Cat. no. 1011/5). Each drug solution was freshly prepared before the experiment. Catechin was firstly dissolved in 96% ethanol to give a stock solution of 5 mM concentration and afterwards diluted to 10 μM concentration with culture medium. Honokiol and curcumin were dissolved in dimethyl sulfoxide (DMSO, Sigma-Aldrich, Cat. no. D8418) to the 5 mM concentration and subsequently diluted to 10 μM (curcumin) and 2 μM (honokiol) concentrations with growth medium. Cinnamon was prepared by dilution in the PBS buffer to the 1000 μg/L concentration and then diluted again to 2 μg/L concentration with culture medium.

### 3.3. MTT assay

The viability of cells was determined using the standard MTT [3-(4,5-dimethylthiazol-2-yl)-2,5-diphenyltetrazolium bromide] assay (MTT, Sigma-Aldrich, PL). The cells were incubated on 96-well plate (Perkin Merkel) in the concentration 10 000 cells/well for 3 hours (37°C) in 0,5 mg/ml MTT solution in PBS buffer (100 μl/well). The absorbance was measured in 0h (directly after the balloon flight), 24h, 48h and 72h incubation after balloon landing at 570 nm (Multiplate reader EnSpire, Perkin Elmer). Acidified isopropanol (100 μL/well, 0.04 M HCl in absolute isopropanol per well) was used to dissolve the formed formazan crystals.

### 3.4. Cell death assay

After landing, the cells were plated on 6-well plates (200 000 cells/well) and incubated for 24 hours in the appropriate drug solution. Following incubation, the culture medium was removed cells were washed with PBS buffer and detached from the surface using trypsin-EDTA solution. Subsequently, they were stained with Annexin V-APC Apoptosis Kit with PI (BioLegend, Cat. no. 640932) and analyzed with FACS Calibur flow cytometer (Becton Dickinson) in order to indicate the percentage ratio of early and late apoptotic and necrotic cells.

### 3.5. Membrane permeabilization assay

Fluorescent dye YO-PRO-1 (Invitrogen, Cat. no. Y3603) was used to evaluate the plasma membrane permeabilization resulting from the low-temperature and radiation cell damage during the balloon flight. The cells were seeded on black 96-well plates (Perkin Elmer) and stained in 1 μM YO-PRO-1 diluted in growth medium for 10 minutes. After 10 min. of incubation and washing with PBS, the intracellular fluorescence of YO-PRO-1 was measured with excitation wavelength of 491 nm and the emission wavelength of 509 nm. Membrane permeabilization assay was evaluated directly after balloon flight (0h) and 24h after landing (24h).

### 3.6. Intracellular ROS generation assay

The level of the reactive oxygen species (ROS) in cells was determined with the DCF (2,7-dichlorofluorescein) assay (Life Technologies, Poland). For experiments, the stock solution of carboxy-H2DCFDA (50 µg/mL in sterile DMSO; Sigma, Poland) was established at the RT in the dark and then diluted in a cell culture medium without FBS. The cells were seeded on black 96-well plates. After washing with PBS the reagent was added to the cell culture to a final concentration of 5 µM and cells were incubated at 37°C in darkness for 30 min. After the incubation, the fluorescence of DCF in wells was measured every 30 minutes for 90 minutes total (with excitation wavelength of 495 nm and the emission wavelength of 530 nm). Intracellular ROS generation assay was evaluated directly after balloon flight (0h) and 24 hours after landing (24h).

### 3.7. Immunocytochemical staining

Immunocytochemical staining was performed using the ABC method in order to investigate the effect of stratospheric conditions and antioxidative drugs on the expression of manganese-dependent superoxide dismutase (SOD2) in SKOV-3 and CHO-K1 cells. After landing, the cells were seeded on 10-well diagnostic microscopic slides (Thermo Fisher Scientific) and incubated for 24 hours in the appropriate drug solution. After incubation, cells were fixed and dehydrated using 4% paraformaldehyde (PFA, Sigma-Aldrich,) for 10 minutes. The, the cells were stained using the EXPOSE Mouse and Rabbit Specific HRP/DAB Detection IHC kit (Abcam, United States; Cat. no. ab80436). The enzyme expression was visualized with the mouse monoclonal antibody anti-SOD2 (Santa Cruz, USA, Cat. no. sc-362300) diluted 1:200 with the PBS buffer. After the overnight incubation with the primary antibody, cells were incubated with the secondary antibody conjugated with horseradish peroxidase (HRP). Next, samples were incubated with a diaminobenzidine-H2O2 mixture in order to visualize the HRP label. Between particular steps the samples were rinsed using 1% Triton X-100 in PBS. The cells were stained with hematoxylin for 3 minutes to visualize nuclei. The immunocytochemical reaction was evaluated with double-blinded method using upright microscope (Olympus BX51, Japan). Then the percentage of stained cells was estimated and the intensity of immunocytochemical reaction evaluated according the scale: (−) negative, (+) weak, (++) moderate and (+++) strong.

### 3.8. Clonogenic assay

Clonogenic assay is a technique allowing for the assessment of cell survival and proliferation following the exposition to the tested compounds. After balloon landing, the cells were plated in appropriate dilutions (150 cells/well) into 6-well-plates at appropriate drug solution. Multi-well-plates were placed in an incubator and left there for 7 days until large colonies (> 1 mm) were formed (50 cells or more). After incubation, growth medium was removed, and the cells were washed with PBS. Fixation and staining of clones was done with a mixture of 0.5% crystal violet in 4% paraformaldehyde (PFA, Sigma-Aldrich, USA) for 10 minutes. Then, the plates were rinsed with water and left to dry at room temperature. Counting of clones was performed the following day.

### 3.9. Neutral comet assay

For detection of DNA damages associated with the exposure to extreme environment during the balloon flight, neutral comet assay method described by Collins (44) was used. Directly after the balloon flight, CHO-K1 and SKOV-3 cells were suspended in freezing medium Bambanker™ (Nippon Genetics, Cat. no. BB01) (1 × 10^5^ cells /50 μL) and frozen for further analysis. After defrosting and centrifugation with PBS, the cells at the concentration 1 × 10^5^/ml were mixed with low temperature melting agarose (Sigma) at ratio 1:10 (v/v) and spread on a slide. Slides were submerged in precooled lytic solution (2.5 M NaCl, 100 mM EDTA, pH 10, 10 mM Tris base and 1 % Triton X-100) at 4°C for 60 min. After lysis and rinsing, slides were equilibrated in TBE solution (40 mM Tris/boric acid, 2 mM EDTA, pH 8,3), electrophoresed at 1,2 V/cm for 15 min and then silver staining was performed (45). For scoring the comet patterns, 200-300 nuclei from each slide were assessed. Additionally, the samples were stained using fluorescent dye Green Dead Cell Stain (Thermo Fisher Scientific, Cat. no. S34860). The images were acquired on a fluorescence microscope (Olympus Nikon). For each measurement, at least 50 cells per sample were counted and the CometScore 2.0 software was used to analyze the comets. The tail DNA percentage was taken as a quantified index of DNA damage.

### 3.10. Confocal microscopy and cell morphology analysis

Confocal laser scanning microscopy (CLMS) was used for the visualization of cell membrane damage and morphology. After the balloon flight, the cells were grown on coverslips for 24 hours. Subsequently, the cells were washed 3 times with PBS and quickly submerged in staining solution CellMask™ Deep Red Plasma Membrane Stains (at a concentration of 0,5 μg/ml dissolved in growth medium; Molecular Probes, Cat. no. C10046, Ex./Em. 649/666 nm) for 10 minutes at 37°C. Next, the staining solution was removed and the coverslip were rinsed with PBS three times. After washing out with PBS, nuclear DNA was stained with DAPI (4,6-diamidino-2-phenylindole; 0.2 μg/ml, Ex./Em. 358/461 nm). In the end the cells were mounted in fluorescence mounting medium (DAKO). For imaging, Olympus FluoView FV1000 confocal laser scanning microscope (Olympus) was used. The images were recorded by employing a Plan-Apochromat 60× oil-immersion objective.

### 3.11. Stratospheric equipment

The balloon was filled with helium and its position was tracked by APRS-supported services: www.habhub.com and www.aprs.fi. The telemetry was emitted through RTTY 70 cm/APRS 2 m signal, supported with GPS/GSM tracker. On-board computer (ATMEGA328P) was equipped with sensors located at the top wall (for UV radiation (ML8511), pressure (BMP085), and temperature (DHT22) measurements), additional UV sensors at the side wall of gondola and accelerometer (MPU6050) inside the box. To compare stratospheric and Earth UV irradiation, the UV ML8511 sensor was used which is designed especially for surface measurement of UV spectra (UVA (280-400 nm) and UVB (260-280 nm)). As it is mentioned in the specification, this sensor has a nominal operation range of −20°C to 70°C suggesting the monitoring of UV would only be valid up to roughly the minute 30 of the flight. This means that some of the random variation seen in Fig 1C may be an artefact of the electronics which is not operating adequately at these lows temperatures. It often happens that the electronic produces too much noise when it goes below certain temperature thresholds and furthermore here is exposed to moisture freezing. However, grounded analyses displayed that the sensor may be used in temperatures below −20 which cause insignificant disturbances of measurements.

### 3.12. Statistics

Statistical significance was determined by unpaired T-Student tests for cytotoxicity tests and two-way analysis of variance (ANOVA). Differences between treated samples and control cells with p values ≤0.05 were assumed to be statistically significant. The results were analyzed with the Microsoft Office Excel 2017 and GraphPad Prism 7.0 software.

## 4. Results

### 4.1. Balloon flight and stratospheric conditions analysis

The biological samples were launched to the stratosphere on 30th of April 2018, from Wrocław, Poland (51°06′23.6″ N 17°03′32″ E) at 11:30 AM. The balloon reached the stratosphere at maximal altitude of 30 298 m. The mission lasted about 2 hours: 90 min of ascent and 25 min of descent and ended at 1:25 PM. The biological samples were collected directly after landing in Sulisław, Poland (52°23′49.9″ N 18°45′52.8″ E) and transported to the laboratory. During the first stage of ascending phase recorded ambient temperature dropped to −22°C. Subsequently, when the balloon reached the ozonosphere, the ambient temperature increased to −2°C. At the highest altitude, the temperature reached the lowest level of −35°C (Fig 3A) and the lowest pressure (1252 Pa) was measured (Fig 3B). Measurements provided by two UV sensors show rapid variations of UV radiation, indicating continuous rotations of the gondola (Fig 3C). Voltage level of 1170 mV correlates with the highest score (11) in the UV Index-exposition scale, which shows extreme exposure to the UV radiation causing immediate damage of unprotected human skin and eyes (46). On the right side of the chart there is a clearly visible period when the parachute was opened, and the gondola was stabilized. In the upper parts of the atmosphere, the UV dose was greater more than two times than the dose correlating with the maximum dose in the UV-Index scale (reaching nearly 2463 mV).

**Fig 3.**
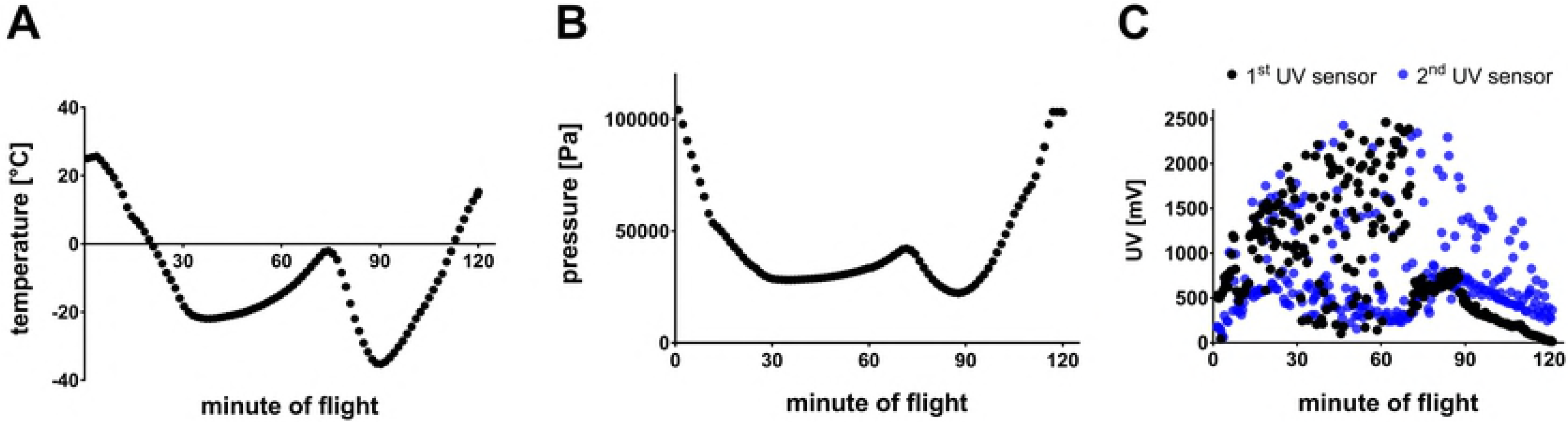
The environmental analysis. (A) The temperature fluctuations [°C] during the balloon flight outside the payload; the sensor was located at the top wall of gondola. (B) The pressure fluctuations [Pa] during the balloon flight, the sensor was located at the top wall of gondola. (C) UV radiation fluctuations [mV] during the balloon flight. The 1^st^ UV sensor was located at the top wall of gondola, the 2^nd^ UV sensor measured UV radiation at the side wall.

### 4.2. Cells viability

Following the flight, we observed a decrease of the mitochondrial activity of CHO-K1 and SKOV-3 cells not protected against radiation and not incubated with antioxidants (Fig 4A, 4C). The presence of antioxidants, except for curcumin, caused increase in the CHO-K1 cells’ viability. The most protective effect of antioxidants was observed in CHO-K1 cells incubated with catechin and honokiol. Studies have shown a slight increase of mitochondrial activity in normal cells after treating with cinnamon. Curcumin was the most lethal for these cells (72h - 25%) (Fig 4A). There was also observed a significant increase in the mitochondrial activity in CHO-K1 cells protected against radiation, especially after catechin and honokiol treatment (Fig 4B).

**Fig 4.**
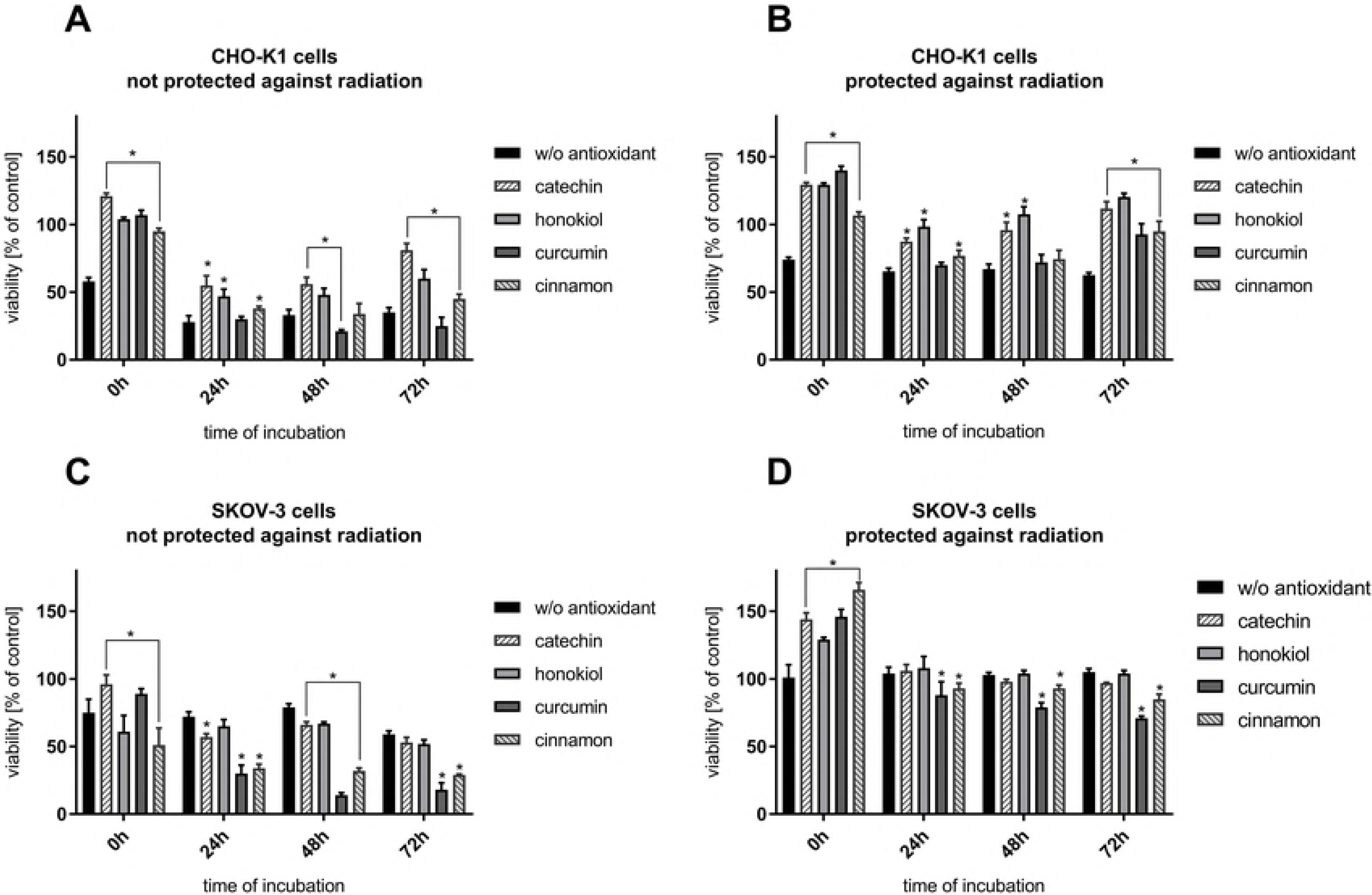
Cells viability verified by MTT assay after the balloon flight. (A) CHO-K1 cells not protected against radiation, (B) CHO-K1 cells protected by the aluminum foil against radiation, (C) SKOV-3 cells not protected against radiation, (D) SKOV-3 cells protected by aluminum foil against radiation (*p ≤ 0.05). Data are presented as the mean percentage relative to control cells, which were not sent into the stratosphere.

The results have shown that SKOV-3 cells not protected against radiation display similar mitochondrial activity after balloon flight - about 70% in comparison to CHO-K1 cells (Fig 4C). Furthermore, it was observed that curcumin and cinnamon were the most lethal for SKOV-3 cells, as their mitochondrial curcumin and cinnamon activity was decreasing repetitively approximately to the 18% and 30%. Other substances used in this experiment exert the less lethal effect on SKOV-3 cell line. The most protective antioxidant for that cells was catechin. However, there was no increase of mitochondrial activity in SKOV-3 cells incubated with honokiol. It was observed that curcumin significantly diminished mitochondrial metabolism in both cell lines.

The mitochondrial activity of cancer cells incubated without antioxidants and protected against radiation remained unchanged (approximately 100%) in all 72 hours (Fig 4D). Directly after the flight we observed the highest mitochondrial activity when the antioxidants were applied. A significant decrease was observed after usage of curcumin – 88% after 24 hours, 79% after 48 hours and 71% after 72 hours. However, a smaller but still significant drop was observed after cells’ incubation with cinnamon. Catechin and honokiol application generated results similar to the control cells and cells without antioxidants.

### 4.3. Intracellular reactive oxygen species (ROS) generation

Directly after the stratospheric flight (0h), a significant decrease in intracellular ROS generation in CHO-K1 cells not protected against radiation in case of pre-treatment with catechin and honokiol was observed (Fig 5A). However, after 24 hours, an increased level of ROS was detected. In case of CHO-K1 not protected against radiation and pretreated with catechin, the lowest and relatively constant level of ROS was detected. Those results confirmed that catechin displays the most valuable protective role from all of the tested compounds.

**Fig 5.**
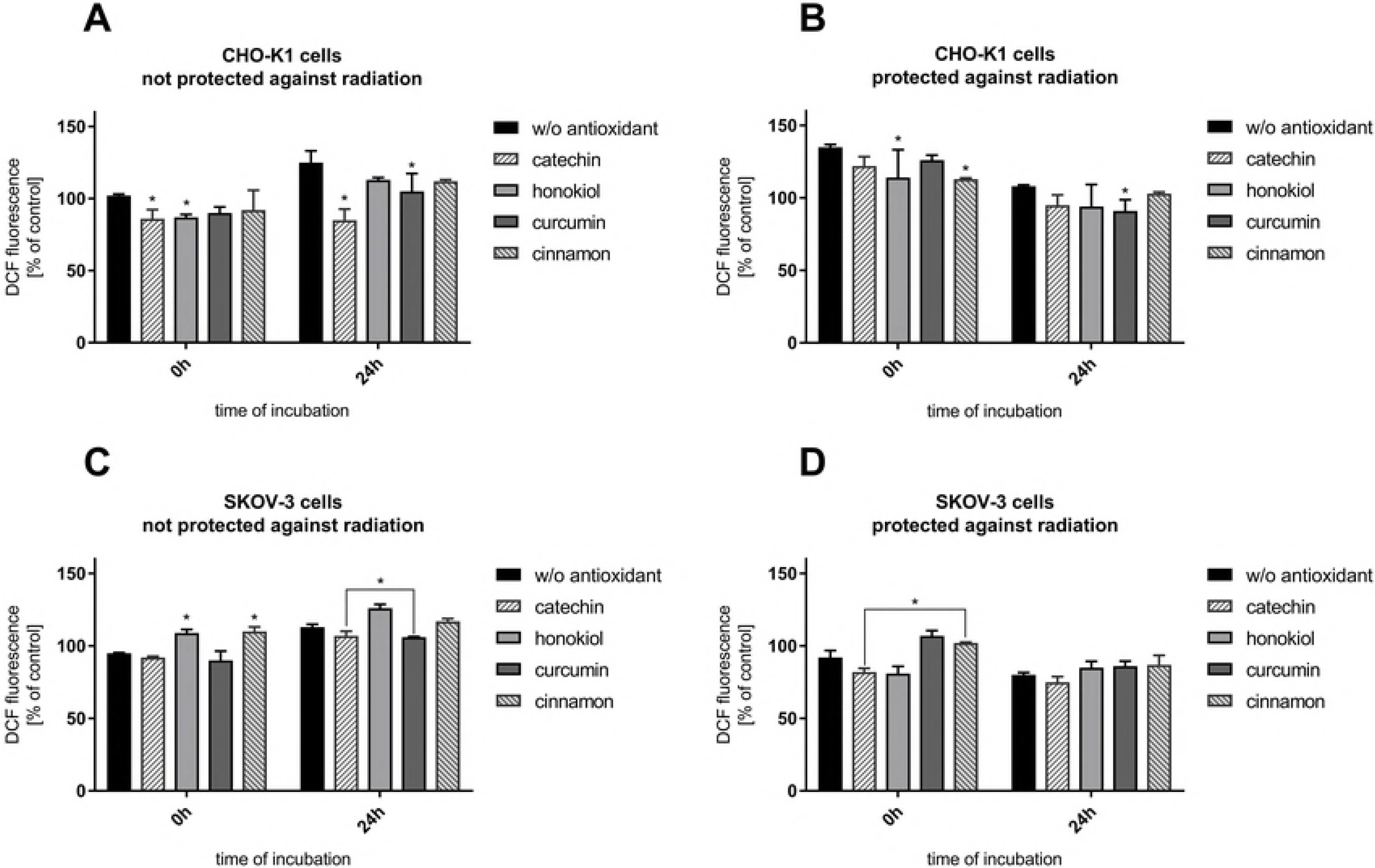
Intracellular reactive oxygen species generation verified by DCF assay after the balloon flight. (A) CHO-K1 cells not protected against radiation, (B) CHO-K1 cells protected by the aluminum foil against radiation, (C) SKOV-3 cells not protected against radiation, (D) SKOV-3 cells protected by aluminum foil against radiation (*p ≤ 0.05). Data are presented as the mean percentage relative to control cells, which were not sent into the stratosphere.

Furthermore, directly after the balloon flight and 24h incubation there was a reduced level of oxidative stress after pretreatment with various antioxidants in CHO-K1 cells protected against radiation measured (Fig 5B). However, it was still similar to the cells incubated in antioxidant-free medium.

Moreover, our studies revealed that in SKOV-3 cells not protected against radiation, the level of oxidative stress after pretreatment with curcumin and catechin is almost the same as in the cells incubated without antioxidant (Fig 5C). Treatment with honokiol and cinnamon resulted in the increase of ROS generation.

In the case of SKOV-3 cells protected against radiation, a higher level of ROS in cells incubated with curcumin and cinnamon was observed (Fig 5D). The results indicated that those compounds could have display antitumor activity by generating reactive oxygen species in cancer cells, which was reflected in our research. After 24 hours, the level of oxidative stress for all the tested compounds and cells without antioxidant was at 80%.

In summary, our results have shown that intracellular ROS generation is related to the exposure to stratospheric conditions, especially radiation, and treatment with the examined antioxidants which act differently in various cells.

### 4.4. Membrane permeabilization

Reduced membrane permeabilization is crucial due to the greater cells protection against harmful radiation and temperature shock. Directly after the stratospheric balloon flight (0h), high membrane permeabilization in CHO-K1 cells not protected against radiation was observed (Fig 6A). After 24 hours the membrane permeabilization for all samples decreased significantly. However, it was observed that the cells after preincubation with catechin have less permeable cell membranes when compared to the other samples. In the case of CHO-K1 cells not protected against radiation, a constant level of cell membrane permeabilization regardless to the type of antioxidant was observed (Fig 6B).

**Fig 6.**
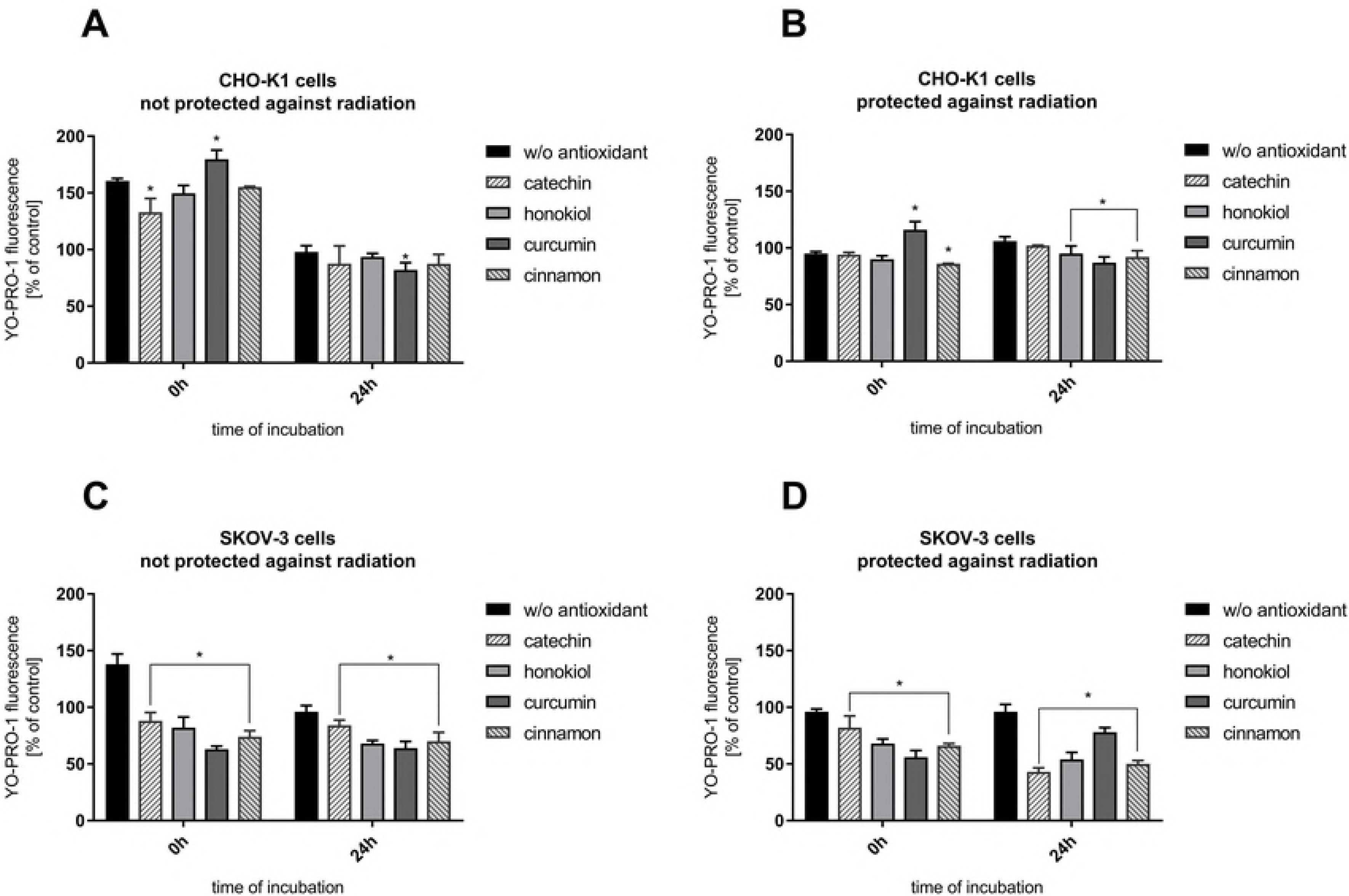
Cell membrane permeabilization determined by YO-PRO-1 fluorescence after the balloon flight. (A) CHO-K1 cells not protected against radiation, (B) CHO-K1 cells protected by the aluminum foil against radiation, (C) SKOV-3 cells protected by aluminum foil against radiation (*p ≤ 0.05). Data are presented as the mean percentage relative to control cells, which were not sent into the stratosphere.

Our studies indicated a decreased membrane permeabilization in SKOV-3 cells not protected against radiation and incubated with antioxidants (Fig 6C). The same dependency was observed in case of SKOV-3 cells protected against UV radiation (Fig 6D). Of all the tested compounds, once again catechin showed the best protective effect due to the reduced permeability of the plasma membrane.

### 4.5. Cell death

Cell death assay revealed different values of apoptotic and necrotic cells after the balloon flight. We noticed the increased percentage ratio of necrotic cells in case of CHO-K1 cells exposed to radiation in comparison to CHO-K1 cells protected against radiation (Fig 7A-B, Tab. 1, Fig 8). Incubation with catechin, honokiol or curcumin resulted in a reduced level of total apoptotic cells, whereas the cinnamon revealed the strongest proapoptotic agent.

**Fig 7.**
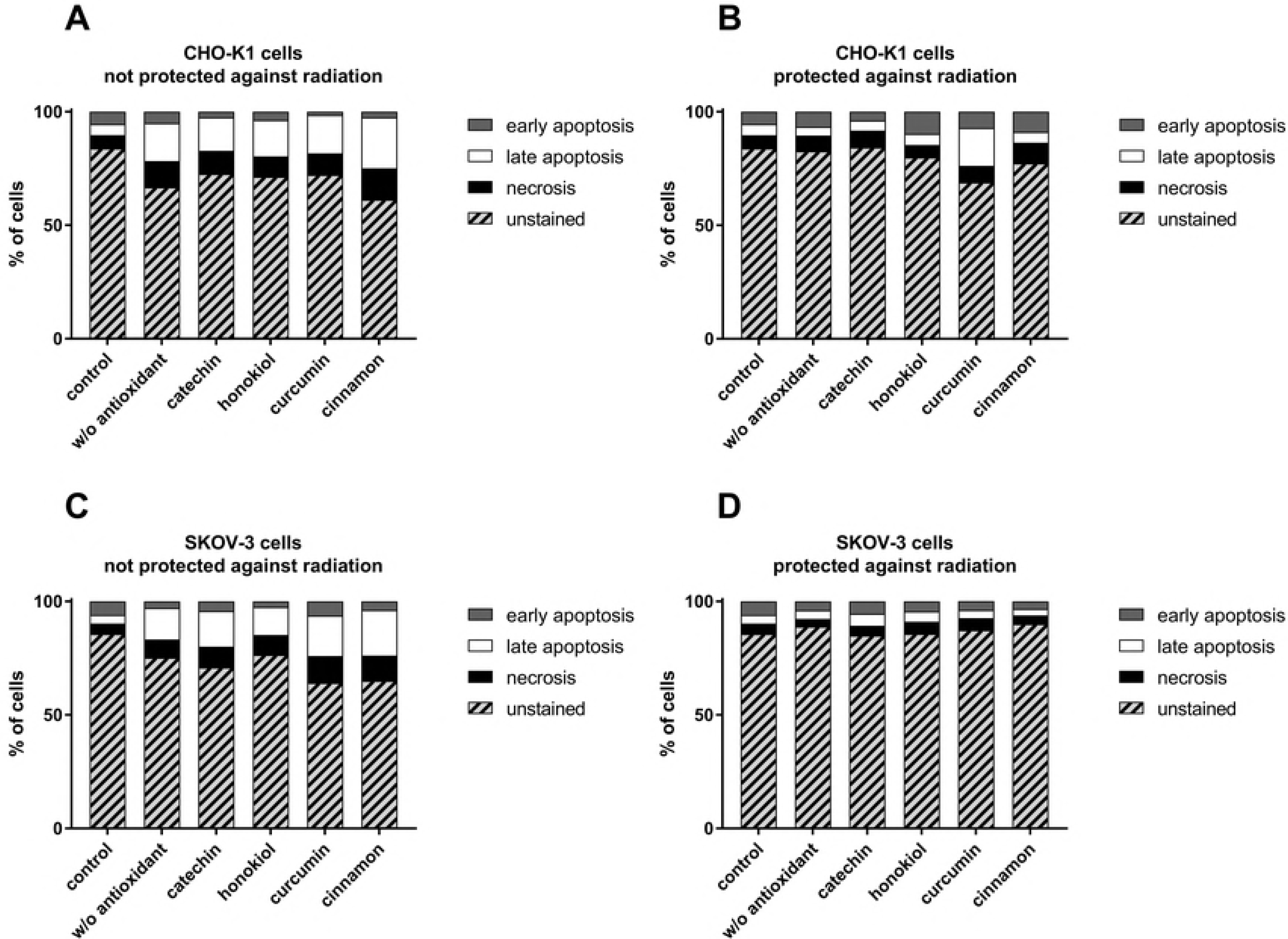
The percentage ratio of undamaged, necrotic, early and late apoptotic cells evaluated by Annexin V-APC/PI staining 24-hour after balloon flight. (A) CHO-K1 cells not protected against radiation, (B) CHO-K1 cells protected by the aluminum foil against radiation, (C) SKOV-3 cells not protected against radiation, (D) SKOV-3 cells protected by aluminum foil against radiation.

**Tab. 1.** Cell death evaluation of CHO-K1 and SKOV-3 cells protected and not protected against radiation, treated with various antioxidants, after 24-hour incubation after balloon flight. Results expressed as percentage of cells.

| CHO-K1 cells | undamaged | necrosis | total apoptosis | SKOV-3 cells | undamaged | necrosis | total apoptosis |
| --- | --- | --- | --- | --- | --- | --- | --- |
| *not protected against radiation* |  |  |  | *not protected against radiation* |  |  |  |
| control | 84 | 6 | 10 | control | 85 | 5 | 10 |
| w/o antioxidant | 67 | 12 | 22 | w/o antioxidant | 75 | 8 | 17 |
| catechin | 73 | 10 | 17 | catechin | 71 | 9 | 20 |
| honokioli | 71 | 9 | 20 | honokioli | 77 | 9 | 15 |
| curcumin | 72 | 10 | 18 | curcumin | 64 | 12 | 24 |
| cinnamon | 61 | 14 | 25 | cinnamon | 65 | 11 | 24 |
| *protected against radiation* |  |  |  | *protected against radiation* |  |  |  |
| control | 84 | 6 | 10 | control | 85 | 5 | 10 |
| w/o antioxidant | 83 | 7 | 10 | w/o antioxidant | 89 | 3 | 8 |
| catechin | 84 | 8 | 8 | catechin | 85 | 4 | 11 |
| honokioli | 80 | 5 | 15 | honokioli | 86 | 5 | 9 |
| curcumin | 69 | 7 | 24 | curcumin | 87 | 5 | 8 |
| cinnamon | 77 | 9 | 14 | cinnamon | 90 | 4 | 6 |

**Fig 8.**
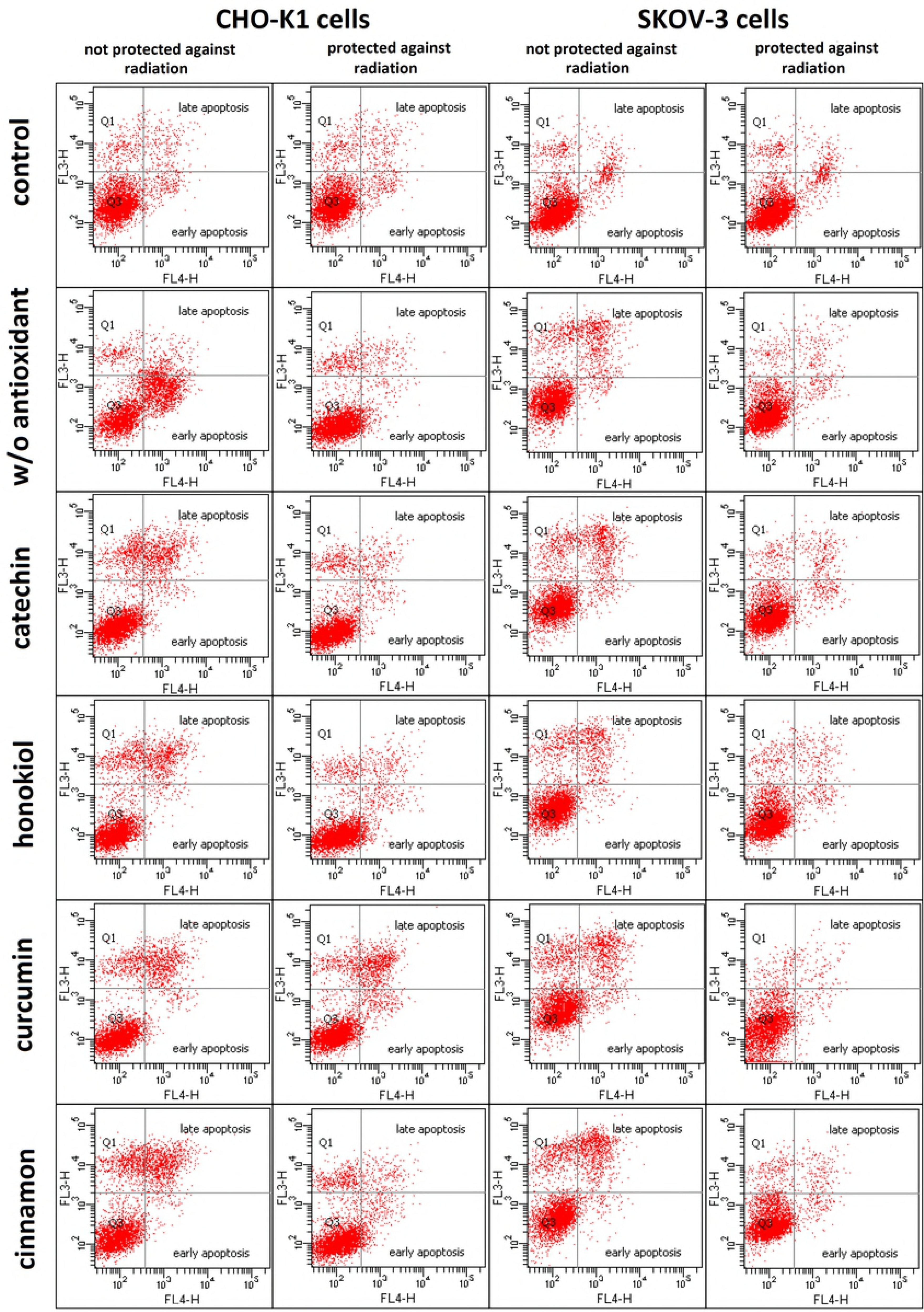
The comparison of early and late apoptotic CHO-K1 and SKOV-3 cells protected by aluminum foil and not protected against radiation, treated with various antioxidants, after 24-hour incubation after the balloon flight. Results are represented as dot plots.

The highest percentage ratio of late apoptotic cells was observed after curcumin treatment of CHO-K1 cells protected against radiation (Fig 7B). Interestingly, incubation with honokiol and cinnamon resulted in the increased level of early apoptotic cells. The lowest level of total apoptotic cells was observed after the catechin treatment.

SKOV-3 cells not protected against radiation showed the increased percentage ratio of apoptotic cells (Fig 7C) in comparison to the cells protected against radiation. The highest level of dead cells was observed after treatment with curcumin and cinnamon, whereas honokiol exhibited antiapoptotic properties.

In case of SKOV-3 cells protected against radiation, similar percentage ratio of apoptotic cells, for the cells both incubated with incubated with and without antioxidants was observed. (Fig 7D). The highest level of necrotic cells was revealed for the cells treated with honokiol and curcumin. However, the reduced percentage ratio of apoptotic cells was detected only for the cells incubated with cinnamon.

### 4.6. SOD2 expression

Immunocytochemical staining method enabled detection of the significant differences in SOD2 expression after the balloon flight in both examined cell lines (Fig 9, Tab. 2). After the stratospheric flight, we observed a weaker expression of the manganese-dependent superoxide dismutase in SKOV-3 cells in comparison to CHO-K1 cells in the presence of the same antioxidant compounds. CHO-K1 cells treated with catechin and cinnamon extract were the most immunoreactive. After the balloon flight only CHO-K1 cells exhibited the increased SOD2 reactivity in comparison to control cells whereas the opposite changes were observed in SKOV-3 cells. The obtained results indicate that the examined antioxidants induced SOD2 activity in normal cells more effectively than in cancer cells.

**Fig 9.**
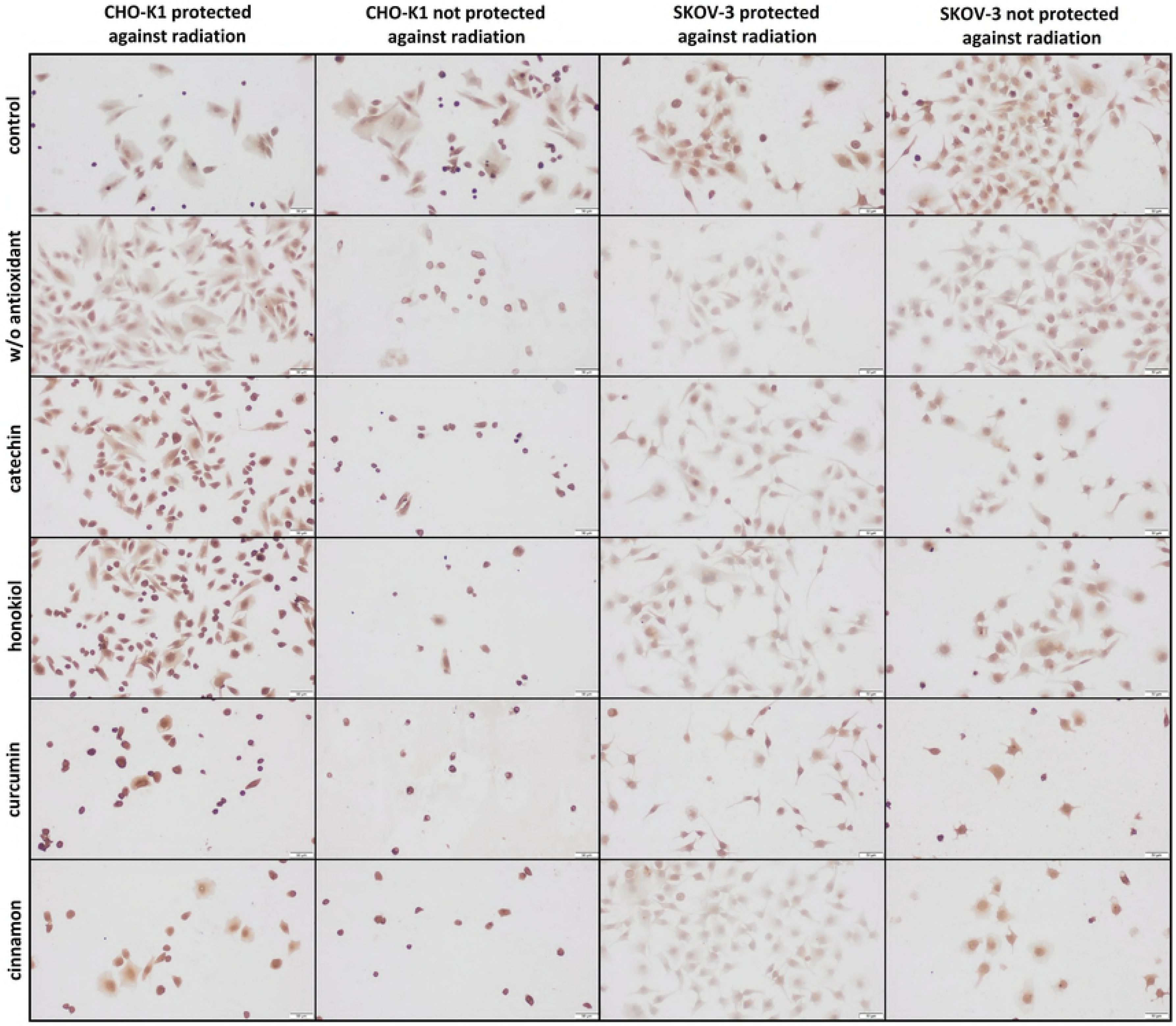
The representative microphotographs of the immunoreactivity of the manganese-dependent superoxide dismutase (SOD2) in CHO-K1 and SKOV-3 cells protected and not protected against radiation.

**Tab. 2.** The immunocytochemical reaction with SOD2 antibody in CHO-K1 and SKOV-3 cell lines after the balloon flight in the presence of various compounds, taking into account the exposure and the protection against radiation (% - percentage of stained cells, RI - Reaction Intensity).

|  | control |  | w/o antioxidant |  | catechin |  | honokioli |  | curcumin |  | cinnamon |  |
| --- | --- | --- | --- | --- | --- | --- | --- | --- | --- | --- | --- | --- |
|  | % | RI | % | RI | % | RI | % | RI | % | RI | % | RI |
| CHO-K1 protected against radiation | 90% | +/++ | 100% | ++ | 95% | ++/<br>+++ | 95% | +++ | 100% | +++ | 100% | ++/<br>+++ |
| CHO-K1 not protected against radiation | 80% | ++ | 85% | +/++ | 90% | +++ | 100% | ++/++<br>+ | 100% | ++/++<br>+ | 100% | +++ |
| SKOV-3 protected against radiation | 95% | ++/++<br>+ | 50% | + | 70% | +/++ | 80% | +/++ | 90% | ++ | 70% | +/++ |
| SKOV-3 not protected against radiation | 95% | ++/++<br>+ | 70% | +/++ | 100% | +/++ | 95% | ++ | 100% | ++ | 100% | ++ |

### 4.7. Clonogenic assay

The clonogenic assay confirmed the protective role of selected antioxidants which enhanced cell survival after exposure to the stratospheric environment (Fig 10, 11). Differences in number of colonies between the protected and unprotected cells indicate the cytotoxic properties of radiation. Furthermore, this assay confirmed that CHO-K1 were more sensitive to high energy particles in the stratosphere. We observed more and bigger colonies in CHO-K1 cells, especially among the protected cells. The least efficient in cell protection was curcumin whereas the highest number of colonies was observed after cinnamon and catechin pretreatment.

**Fig 10.**
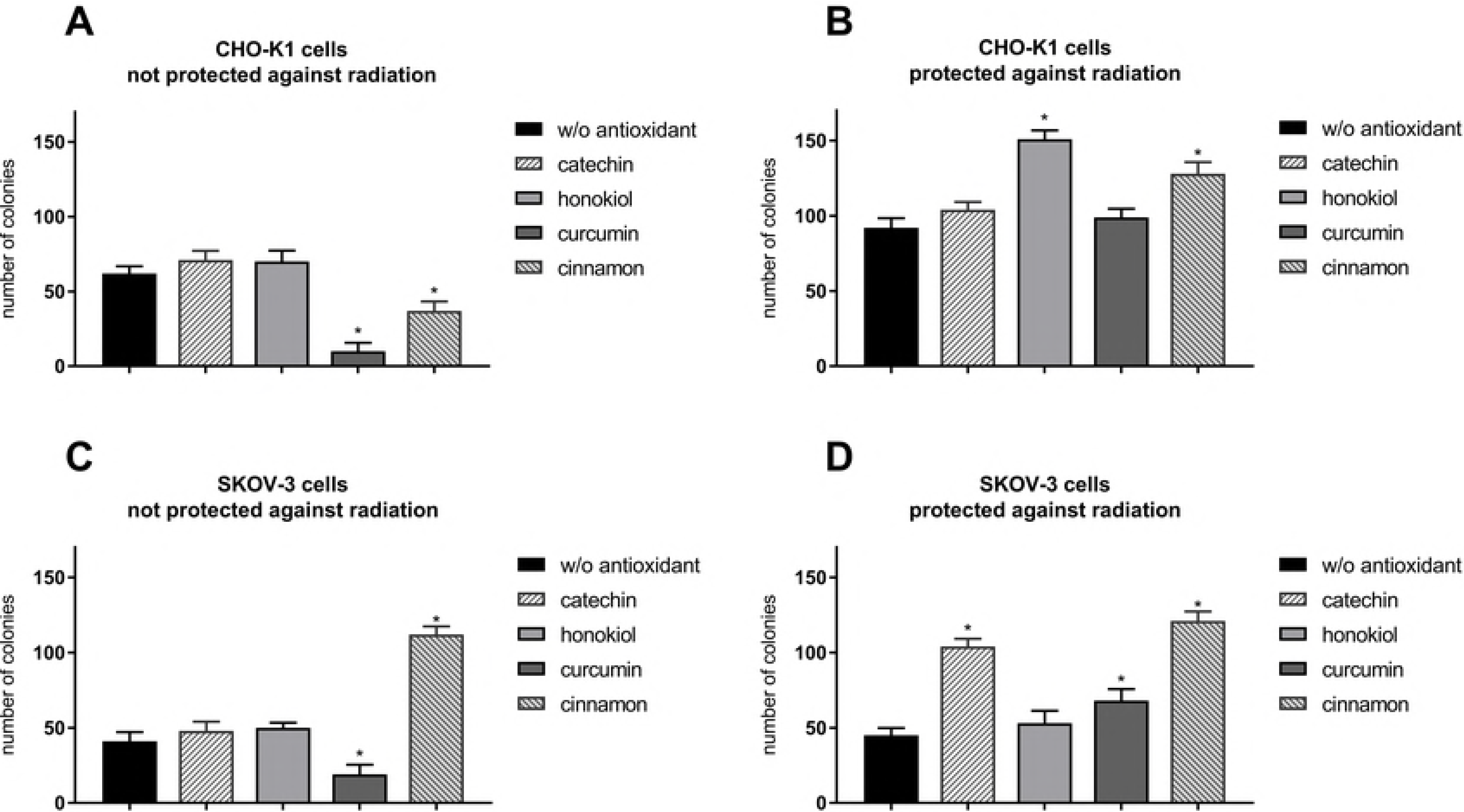
Number of colonies after 7-day incubation. (A) CHO-K1 cells not protected against radiation, (B) CHO-K1 cells protected by aluminum foil against radiation, (C) SKOV-3 cells not protected against radiation, (D) SKOV-3 cells protected against radiation (*p ≤ 0.05). Data are presented as the mean percentage relative to control cells, which were not sent into the stratosphere.

**Fig 11.**
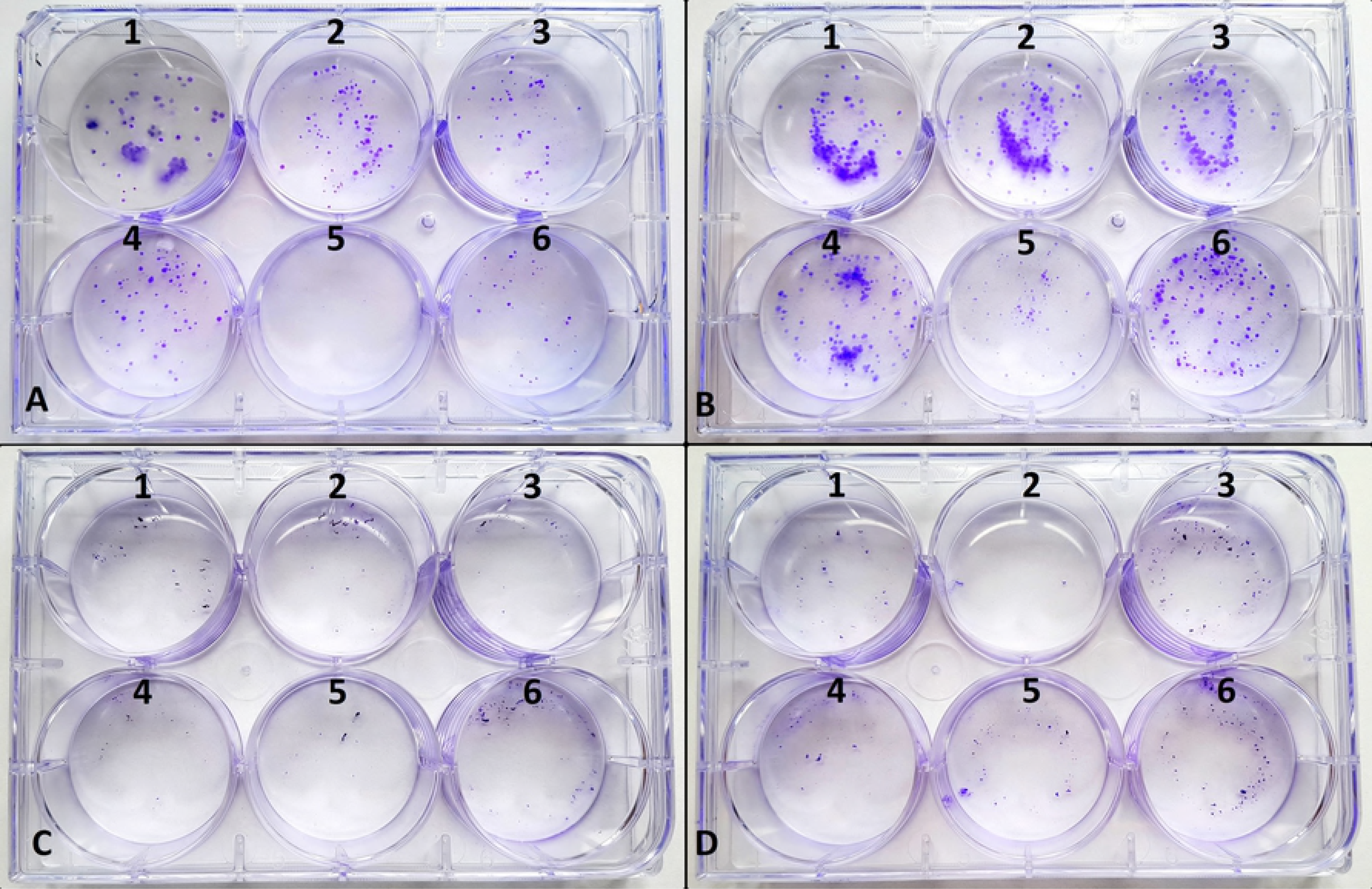
Colony formation in clonogenic assay after 7 day incubation. (A) CHO-K1 cells not protected against radiation, (B) CHO-K1 cells protected against radiation, (C) SKOV-3 cells not protected against radiation, (D) SKOV-3 cells protected against radiation, 1 - control cells, 2 - w/o antioxidant, 3 - catechin, 4 - honokiol, 5 - curcumin, 6 - cinnamon.

Comparison of MTT, cell death and clonogenic assay for SKOV-3 cells not protected against radiation and treated with cinnamon revealed a decrease of cell viability after the balloon flight, however, proliferation ability was not affected. It suggests that cinnamon first, may cause increased cell death after the flight and then promote cells’ multiplying. We did not observe this effect for curcumin treatment.

### 4.8. DNA damage in the stratosphere

In neutral comet assay we were able to distinguish four types of comets. Figure 12 shows the detailed categorization method used for the evaluation of our samples that allows us to divide the observed nuclei into three groups: undamaged, apoptotic and intermediately damaged (Tab. 3). Comparison of CHO-K1 and SKOV-3 cells not protected against radiation revealed protective role of antioxidants in DNA damage among CHO-K1 cells, however, curcumin and cinnamon promoted DNA destruction in SKOV-3 cells. In the cells protected against radiation antioxidants acted differently. In the normal cells we observed more undamaged cells after preincubation with catechin and honokiol, whereas in the cancer cells cinnamon additionally reduced the percentage ratio of cells with DNA damages. These results suggest that the protective role of antioxidants in DNA damages depends on the presence of radiation resulting in various activity of these substances in cells.

**Fig 12.**
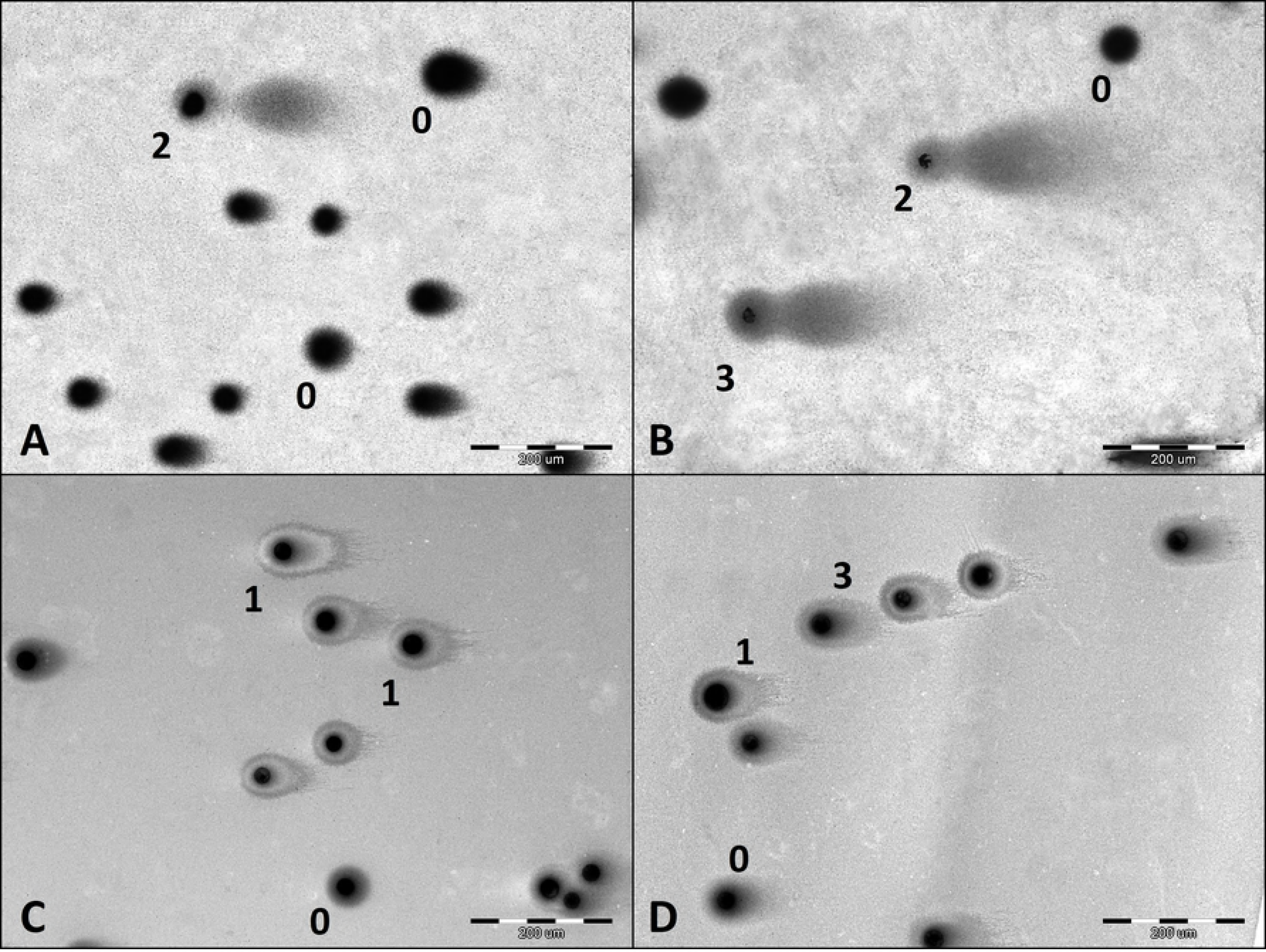
Different types of DNA damage visualized by comet assay. (A) comet type 0 and 2; (B) comet type 0, 2 and 3; (C) comet type 0 and 1; (D) comet type 0, 1 and 3. Type 0 - size of the head the normal nucleus size, tail absent; type 1 - size of the head the normal nucleus size, tail size less than normal nucleus size, type 2 - size of the head less than half size of normal nucleus, tail size more than 2 times the normal nucleus size, type 3 - size of the head the normal nucleus size, tail size about 2 times the normal nucleus size.

**Tab. 3.** DNA damage evaluation of CHO-K1 and SKOV-3 cells protected and not protected against radiation, treated with various antioxidants, directly after balloon flight. Results expressed as percentage of cells.

| CHO-K1 cells | undamaged | apoptotic damaged | intermediately damaged | SKOV-3 cells | undamaged | apoptotic damaged | intermediately damaged |
| --- | --- | --- | --- | --- | --- | --- | --- |
| *not protected against radiation* |  |  |  | *not protected against radiation* |  |  |  |
| w/o antioxidant | 79 | 9 | 12 | w/o antioxidant | 71 | 1 | 28 |
| catechin | 84 | 3 | 13 | catechin | 82 | 1 | 18 |
| honokiol | 84 | 1 | 15 | honokiol | 83 | 1 | 16 |
| curcumin | 91 | 1 | 9 | curcumin | 64 | 0 | 36 |
| cinnamon | 83 | 7 | 10 | cinnamon | 61 | 1 | 38 |
| *protected against radiation* |  |  |  | *protected against radiation* |  |  |  |
| w/o antioxidant | 90 | 2 | 8 | w/o antioxidant | 54 | 1 | 45 |
| catechin | 91 | 0 | 9 | catechin | 74 | 0 | 25 |
| honokiol | 93 | 2 | 5 | honokiol | 83 | 1 | 16 |
| curcumin | 83 | 0 | 17 | curcumin | 52 | 0 | 48 |
| cinnamon | 89 | 2 | 8 | cinnamon | 70 | 1 | 29 |

Furthermore, using CometScore software we were able to perform the analysis of the pictures of comets and determine the percentage ratio of the damaged DNA accompanying the flight (Fig 13). Our studies showed that cancer cells were more vulnerable to UV radiation exposure causing an increase of the presence of DNA in tail. The most active compound preventing DNA damage was cinnamon extract, both for normal and cancer cells. In the case of protected cells, honokiol and curcumin enabled the most efficient preservation.

**Fig 13.**
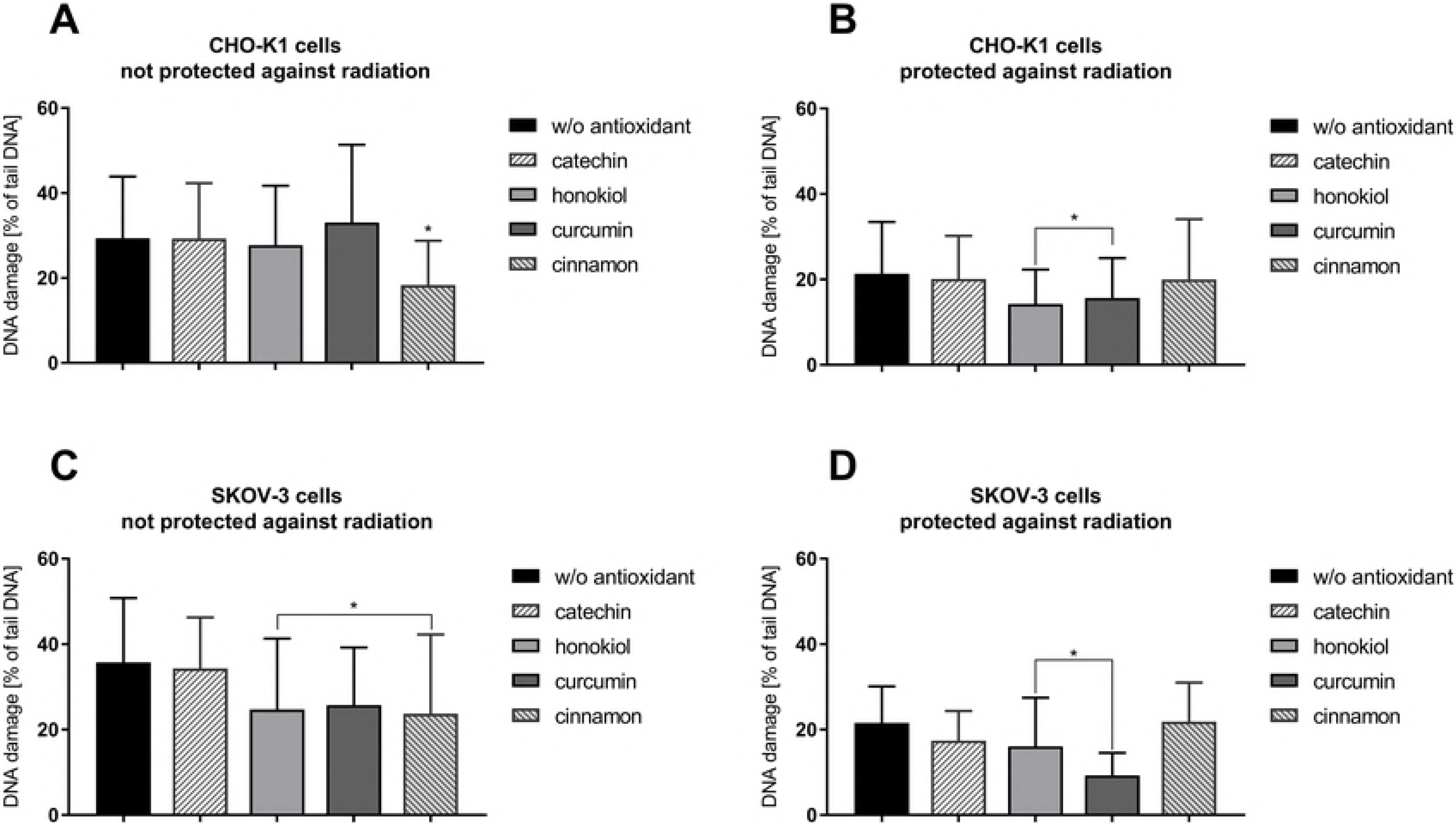
DNA damage evaluation of CHO-K1 and SKOV-3 cells protected and not protected against radiation, treated with various antioxidants, directly after balloon flight. (A) CHO-K1 cells not protected against radiation, (B) CHO-K1 cells protected against radiation, (C) SKOV-3 cells not protected against radiation, (D) SKOV-3 cells protected against radiation (*p ≤ 0.05). Results expressed as percentage of damaged DNA presented in tail of comet.

### 4.9. Confocal microscopy and cells’ morphology analysis

Fluorescence staining revealed slightly differences in the cell morphology after the balloon flight in both examined cell lines (Fig 14). After the stratospheric flight, we observed more slender spikes in both cell lines. CHO-K1 and SKOV-3 cells treated with honokiol, curcumin and cinnamon extract were morphologically similar to the control cells. The cells exposed to radiation were enlarged, especially CHO-K1 cells incubated without antioxidant and protected against radiation. Some nuclei were fragmented, in particular in the cells not protected against radiation. However, the incubation with antioxidant resulted in an increased percentage ratio of cells with condensed nuclei with many nucleoli.

**Fig 14.**
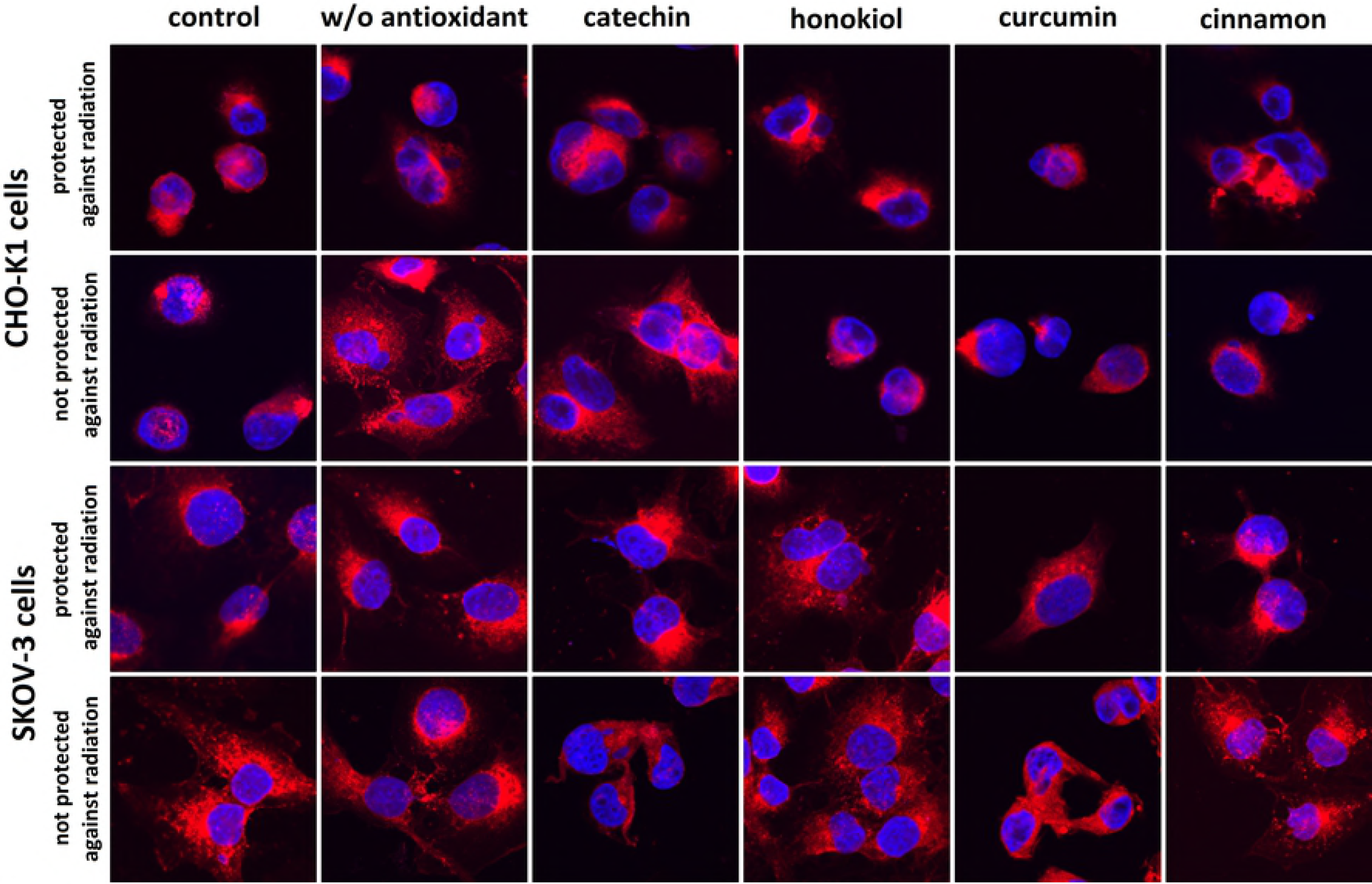
The representative photographs of the morphology of CHO-K1 and SKOV-3 cells protected and not protected against radiation stained with CellMask™ Deep Red (cell membranes) and DAPI (nuclei).

## 5. Discussion

During the balloon flight the cell samples were exposed to several fluctuating stressful factors, namely radiation, temperature, pressure and overload. However, as long as the exposure to these factors does not reach extreme levels, most of the cells are still able to survive due to their natural capability of activating a variety of specific defensive pathways. The stress response could be dramatically different among various types of cells, notably normal and cancer cells. Therefore, the application of meticulously selected compounds could either alleviate or exacerbate the consequences of the exposure to rapidly varying temperature, pressure and UV conditions during stratospheric balloon campaign - depending on their various properties and type of cells - hence influencing not only the survival but also recovery of cells after the experiment. Natural medicines, such as catechin, curcumin, cinnamon extract and honokiol have been recently gaining much attention as therapeutic compounds for cancer prevention and treatment. Our study represents the first attempt at discovering compounds exhibiting protective properties in space by launching the human normal and cancer cells into the stratosphere on a meteorological balloon.

There is a huge ambiguity among the available data concerning quantification of the absorbed dose of cosmic rays depending on the length of the flight and height reached during the meteorological balloon mission. The intensity of the radiation varies depending on parameters such as location and time unit, latitude and longitude, ozone depth and solar activity (47). The average dose of ionizing radiation reaching the atmosphere is estimated at around 2-20 μSv (48). In the stratosphere, 30 000 meters above the Earth’s surface, the UV radiation is much higher than on the sea level (6). The lower the height, the thicker the protective layer of the atmosphere and the higher the radiation exposure (49). During the flight the cells were deprived of the protective effect of the ozonosphere and troposphere (which naturally protect Earth against radiation), resulting in decreased cell viability after the flight.

During the experiment half of the samples was covered by aluminum foil. Therefore, we were able to evaluate the effect of radiation (mainly UV and high energy particles such as α, β^−^) on examined cells in the presence of various compounds. We observed an increased mitochondrial activity and decreased level of apoptotic cells after pretreatment with various antioxidants, especially among CHO-K1 cells. Catechin was the most efficient in radioprotection of normal cells, whereas honokiol appeared to be the most protective for cancer cells. These results were reflected by intracellular ROS measurements and comet assay. We did not observe significant differences in DCF fluorescent signal between SKOV-3 cells protected and not protected against radiation. However, in MTT and cell death assay and immunocytochemical staining the results were altered. It could indicate that radiation-related death in SKOV-3 cells was not strictly related to the oxidative stress and caused direct cell damage.

According to the time of cell death after the irradiation, two types of the apoptosis can be distinguished: fast apoptosis occurring straight after G2 arrest and late apoptosis, when cell death is preceded by one or more cell divisions. Widely understood radiosensitivity has been demonstrated to vary among different cell types or even populations (10). For instance, radiosensitive cells such as thymocytes (50) or lymphocytes (50) tend to undergo fast apoptosis, whereas Yanagihara et al. (1995) described human gastric epithelial tumor cells which exhibited delayed radiation-induced apoptosis scarcely 12h after irradiation, reaching maximum from 72 to 96h (51). We observed a higher percentage ratio of apoptotic cells among normal cells (CHO-K1) as compared to cancer cells (SKOV-3). Our studies suggest that normal cells are more radiosensitive to fast apoptosis than tumorous cells, which resulted in earlier and more intensive apoptosis among CHO-K1 cells. The significant differences were also shown in response to different antioxidants of each cell type, which should be highlighted as particularly interesting in case of curcumin, which acted as light-sensitizer and induced cell death more effectively in SKOV-3. Similar results were observed after treatment with catechin, which promoted apoptosis more frequently in SKOV-3 than CHO-K1. Furthermore, we noticed less and smaller SKOV-3 colonies in comparison to CHO-K1 in clonogenic assay that indicated a delayed radiation-induced death of cancer cells. Additionally, the performed tests highlighted the considerable impact of honokiol and catechin on recovery of the cells after the flight, proving cytoprotective role of antioxidant after radiation exposure.

Radiation-related DNA strand breaks, confirmed in comet assay, have been demonstrated to initiate the expression of p53 (52), which promotes cytochrome C release from the mitochondria and activation of caspase cascade leading to apoptosis (54). SOD2 is the essential mitochondrial antioxidant enzyme which plays a crucial role in protection against radiation in cells (55,56) by scavenging the free radical superoxide (57). Overexpression of SOD2 stabilizes mitochondrial membrane and protect complexes I and III of the respiratory chain from radiation-induced damages (58), decreases the release of cytochrome C, resulting in the apoptosis blockage (59). Due to the intranuclear localization of *SOD2* gene, SOD2 is synthesized in the cytoplasm and subsequently imported into the mitochondria via the mitochondrial protein influx (MPI), where a manganese ion is combined with SOD2 to make active enzyme (60). The research conducted on mammalian cells has demonstrated a decrease of the mitochondrial protein import through MPI caused by radiation which leads to the accumulation of precursor proteins outside mitochondria degraded by proteasomes (61) and the reduced number of mitochondrial proteins. Furthermore, Azzam et al. (2012) revealed that MPI proteins may be damaged by irradiation, resulting in deficient protein import (62). Accordingly, mitochondria play a critical role in radioprotection by affecting SOD2 activity through Cdk1-p53-mediated SOD2 regulation. Transcriptional factors promoted by reduction of ROS production induce transcription of SOD2-regulated genes causing adaptive response to the oxidative stress. Thus, medicines that increase SOD2 activity enhance mitochondrial membrane stability and block the activation of apoptosis. Natural derivatives used in our study have been displayed as radioprotective substances (22,63,64) promoting SOD2 activity ((65–68)). Our studies revealed a significantly increased amount of SOD2 in in normal cells treated with the analyzed antioxidants whereas we observed less viable cancer cells and increased ROS generation among cancer cells treated with these compounds. Curcumin occurred to be the most efficient radioprotective substance in normal cells while it displayed radio- and photosensitive activity in SKOV-3 cells.

According to the data recorded by on-board sensors, the temperature varied between −36°C and 20°C. Lower cultivation temperature is contributed to a decreased cellular metabolism (69), projecting on prolonged preservation of cell viability. However, moderately cold temperature conditions (−60 to −15°C) remain the main limiting factor affecting the cell viability (70). During the flight, the biological samples were exposed to this stressful temperature zone for 2 hours and experienced repeated cycle of chilling and warming while the balloon was ascending and descending (Fig 3): first when temperature dropped down to −21°C, second when it raised to −2°C and third when it dropped down to −36°C before raising to 20°C in the landing area. However, the air density in the stratosphere is about 1000 times lower than on sea level and because of that the temperature exchange between the biological samples and air was limited, resulting in the relative temperature stability of cells. Along with extensive structural damage of the cell, freezing and thawing cycles are associated with oxidative stress and consequently mitochondrial disturbance, cell membrane permeabilization or DNA damage (71), finally leading to death either on apoptotic or necrotic pathway (72,73), which was reflected in our research. The biological samples after the balloon flight were characterized with increase of ROS generation and percentage ratio of dead cells; however, treatment with selected antioxidants limited these processes. We speculate that antioxidants could have reorganized the chemical structure of cell membrane resulting in its increased stability in lower temperature and decreased permeabilization, which was measured in YO-PRO-1 assay directly after the balloon flight and highlighted in CLSM analysis. In this process catechin was the most efficient in CHO-K1 cells. However, we noticed that membrane permeability increased after incubation with curcumin in CHO-K1 cells not protected against radiation after the flight while it was strongly reduced in SKOV-3 cells exposed to the stratospheric conditions.

A number of studies suggest that cryopreservation and low temperature affect the cell sensitivity to radiation (74). According to this, the data concerning utilization of frozen Chinese-hamster-fibroblasts revealed the decrease in X-ray-induced damage of cells stored at (−196)°C in comparison to cells irradiated at room temperature (75). Information from cryocrystallography demonstrates that very low temperatures (−173°C) stop the diffusion of free radicals caused by irradiation which leads to less harmful cell damages induced by X rays (76). Due to this fact, the increased survival of cells affected by freezing may be associated with reduced oxidative stress and damage at low temperature (74). Altogether, the data suggest that freezing can protect cells from radiation-induced damage and apoptosis, affecting cell viability after the flight.

There is a huge ambiguity between different cell types in their response to the temperature-related stress. Our experiment confirms that mechanisms of cellular responses to oxidative stress are strongly associated with the type of a cell. In general, treatment with antioxidants resulted in increased viability in normal cells whereas oxidative stress was intensified in cancer cells. However, the SOD2 expression was higher in normal cells. It shows that molecular antioxidative mechanism of different antioxidants varies in diverse types of cells: the genomic mechanism associated with the increased expression of antioxidative enzymes is predominant in normal cells whereas in cancer cells the antioxidants and their metabolites work as scavengers of free radicals and do not induce intensively the expression of protective proteins.

It is worth noting that the atmospheric pressure during the flight decreased to 1hPa. However, it is not possible to draw indisputable conclusions about the influence of the pressure on the samples due to a fact that in our research the cells were surrounded by the layer of the liquid medium, which remained frozen after the temperature drop. Despite its shortcomings, this method still provides exposure for the decreased pressure to a relatively large extent, which probably influenced the cells homeostasis and hence, the presented data. Notably, among available literature low-pressure cultivating have been proven to influence homeostasis, morphology and proliferation of mammalian cells (77), which become more rounded and less spindle-shaped and show numerous blebs around their margins. Mitochondria and nucleus take more rounded shape due to edema and cell membrane creating bubbles causing cell death (78).

Remarkably, stratospheric balloon flights are accompanied with overloads. Accelerometer in the gondola shows that ascending and descending velocity is not permanent and because of that the biological samples were exposed to overloads. Numerous studies have investigated the effect of overloads on cells. For example, when animal cells are cultured under 10g (10g means 10 times higher force than Earth gravity), proliferation is increased by 20-30%, glucose consumption is reduced, cell migration is inhibited by high overloads. At gravitational stress, cell may move to other metabolic pathways (79). On the other hand, overloads work as a natural antioxidant and reduce lipid membrane peroxidation (LPO). Induction of the passive mechanisms of biomembrane protection associated with changes in the phase status of the membrane is the most plausible explanation for the phenomenon being observed (80). In our experiment we noted changes in membrane permeability, which could have been affected by overloads. Zhan et al. (1999) revealed a protective effect of green tea polyphenols on 10g stress in rats (81) and our studies highlighted the value of natural antioxidants as protective agents in the gravitational stress. Furthermore, altered gravity deeply influences numerous biological processes in living organisms. Changes in gravitational values affect cell survival, development, and spatial organization. In addition, the indirect effects of altered gravity, such as those associated with hydrostatic pressure and fluid shear, strongly affect both *in vitro* and *in vivo* systems (82). Hence, in order to understand these phenomena, further research is necessary.

### 5.1. Conclusions

In our study, were analyzed changes in the functions of normal and cancer cells that occurred due to exposure to high radiation, overloads as well as low temperature and pressure during stratospheric flight. Our work has led us to conclude that the application of the carefully selected medicines enables us to manipulate cellular stress response depending on the type of cells. Altogether, these findings suggest that honokiol and catechin have the best protective effect on the normal cells, whereas curcumin and cinnamon act as radio- and light-sensitizers increasing the percentage of apoptotic cancer cells and DNA damage. The results constitute a significant step towards the investigation of possible strategies for the cell protection in space environment and provide new insights into the application of the examined compounds for the prevention and treatment of cancer. We believe that our research will remain valuable for resolving the difficulty of the human and biological material protection in space. Because of its relatively low costs, our approach remains economic alternative for simulated subcosmic conditions conducted in the laboratory, which requires far more expensive, specialized measurements.

## 6. Acknowledgments

The work was created as part of the activity of the Student Research Group "Biology of Cancer Cell" at the Wroclaw Medical University (SKN No. K 148). The research was supported partially by the Statutory Funds of Wroclaw Medical University no ST.E130.16.060 (manager prof. J. Zalewski), funds of Wroclaw University of Science and Technology and by “Najlepsi z Najlepszych 3.0” program of Polish Ministry of Science and Higher Education. We would like to thank Ms. E. Przydatek for help with the correction of the English language in the manuscript, Mr. J. Skórniak and Mr. W. Tarnowski for organization the balloon flight and transport of the biological samples.

